# Population coding of strategic variables during foraging in freely-moving macaques

**DOI:** 10.1101/811992

**Authors:** Neda Shahidi, Arun Parajuli, Melissa Franch, Paul Schrater, Anthony Wright, Xaq Pitkow, Valentin Dragoi

## Abstract

Until now it has been difficult to examine the neural bases of foraging in naturalistic environments because previous approaches have relied on restrained animals performing trial-based foraging tasks. Here, we allowed unrestrained monkeys to freely interact with concurrent reward options while we wirelessly recorded population activity in dorsolateral prefrontal cortex (dlPFC). The animals decided when and where to forage, based on whether their prediction of reward was fulfilled or violated. This prediction was not solely based on a history of reward delivery, but also on the understanding that waiting longer improves the chance of reward. The decoded reward dynamics were continuously represented in a subspace of the high-dimensional population activity, and predicted animal’s subsequent choice better than the true experimental variables and as well as the raw neural activity. Our results indicate that monkeys’ foraging strategy is based on a cortical model of reward dynamics as animals freely explore their environment.

---

While foraging, animals must continuously make decisions about where to search for food and when to move between possible food sources. To survive in environments with sparse resources, animals forage more effectively if they can predict future outcomes before they execute costly actions such as relocation^1–3^. Two major limitations of past neuroscience studies of foraging have impeded our understanding of this natural behavior. Specifically, trial-based tasks are unable to expose the continuous decision-making process during food search and selection, and experiments conducted with bodily restraints may substantially distort behavior and its underlying natural computations.

Existing foraging theories based on decades of behavioral experiments in various species revolve around the idea of the matching law^4^. This law states that an animal dedicates time or effort to an option in proportion to its value, which is estimated from the history of reward delivery. Tracking reward history on environmentally relevant time-scales enables the animal to detect changes in the reward rate^5,6^. We often quantify how an animal adapts to reward rates by tracking how often it chooses each available option. However, a neglected aspect of adaptive behavior is that the animals adjust their response rate, meaning that they choose ‘when’ to forage in addition to ‘where’ to forage. Choosing the response rate systematically is particularly efficient when the time of choice predicts the chance of receiving a reward, that is, in nature as well as in many foraging studies^4–10^. For a restrained animal engaged in a trial-based foraging task, ‘when’ to choose is distorted by the trial structure, while ‘where’ to choose is distorted by a confined spatial distribution of reward. To examine the time to act, one might run a trial-free task in classical experimental settings in which the animals are chair-seated. However, the freedom to choose ‘when to act’ based on the experimental task is likely distorted for restrained animals by their sense of urgency to finish a session and end the restraints.

Additionally, examining foraging in trial-based tasks makes it difficult to examine the neural bases of the continuous decisions the animal would make freely about when to engage with the task and when to move between food locations.

Restraining animals to record neural activity can cause other inefficiencies in animal behavior^11,12^. The consequences of physical restraints may be especially dramatic on food-seeking behavior because animals use head and body movements to gather information from their environment for foraging^13,14^. Furthermore, the dynamics of neural populations have been shown to differ when the animals aim for targets that are far from their immediate reach^15^.

Here, we address limitations of previous studies by designing a freely moving foraging task, allowing animals to continuously interact with the task and explore a wide range of reward expectancies. We discovered that unrestrained animals do not simply follow the reward flow but predict the instantaneous chance of the next reward and use this estimate to make subsequent choices. We hypothesized that the subjective reward prediction can be decoded from the population activity in dorsolateral prefrontal cortex (dlPFC), an area where neural activity encode reward^10,16,17^ and are related to memory^18^ and motor planning^19^. For decades, recording electrophysiological activity from unrestrained animals has been desired^20^ but technologically challenging. We overcame technical challenges by performing wireless recordings from chronically implanted electrode arrays^21,22^ while designing an experimental setup for the effective transmission of a low power electromagnetic signal (see Methods) ^23,24^.

## RESULTS

Monkeys (*n*=2) were exposed to two concurrent reward sources on a variable interval schedule^7^. We made it costly for the animal to switch between reward sources by placing them 120 cm apart (Fig. 1A, *left*). Animals freely interacted with the task equipment, and we did not impose a trial structure or a narrow response window (see Methods). A multi-electrode Utah array was chronically implanted in dorsolateral prefrontal cortex (dlPFC; Fig. S1) and measured spiking activity was collected using a light-weight, energy-efficient wireless device (Fig 1a, *right* and Fig. 1b).^22^

**Figure 1.**
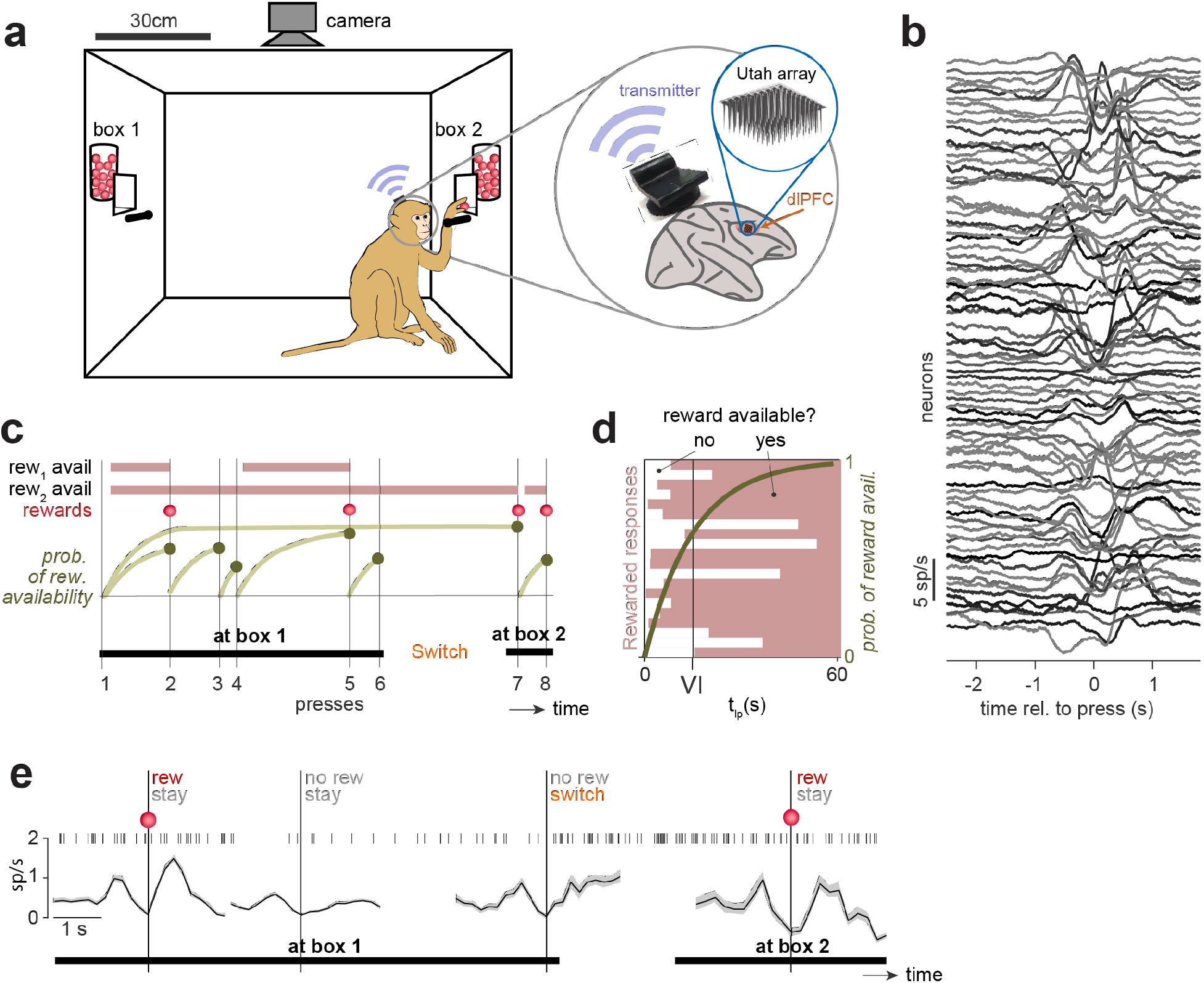
Foraging in freely-moving monkeys while population activity in prefrontal cortex is recorded wirelessly. **(a)** *Left*: Schematic of the experimental setup with two reward boxes, two buttons, and an overhead camera. *Right*: Location of the Utah array in dlPFC (area 46) and wireless transmitter. **(b)** Press-averaged firing rates of 80 single and multi-units recorded simultaneously. **(c)** Illustration of task dynamics with 8 hypothetical presses (vertical lines) in the concurrent variable-interval foraging task. In this illustration, the monkey responds six times on box 1, then switches to box 2 and responds twice. Therefore, press 6 is considered a press with a *switch* choice. The first two rows show the independent telegraph processes determining the reward availability at boxes 1 and 2. In the example shown, press numbers 2, 5, 7, and 8 were rewarded (third row, red). The time dependence of probability of reward availability is shown in the fourth row (see panel d for a different representation). **(d)** The monotonically increasing relationship between the time of the rewards (waiting time) and their preceding presses (shown as pink bars in panel c) and the probability of reward availability (see methods; Fig. S2), shown as a horizontal histogram. **(e)** The spike train of one example neuron on the timescale of four consecutive presses showing a variety of events (top row). Event-locked average firing rates of the same neuron are shown in the bottom row, for conditions with reward/no reward and stay/switch choices. For ease of visualization this example used a neuron with a relatively low firing rate compared to others in the population (compare to panel b).

Rewards on both sides (box 1 and box 2) became available at exponentially distributed random times after the animal obtained a previous reward. The reward availability was hidden from the monkey. Once becoming available, each reward remained available until the animal pressed a button, at which time the reward was delivered (Fig. 1c). The distribution of waiting times before reward became available could have different mean times or ‘schedules’ for each side (*i.e*., constant hazard rates; Fig. 1c). Schedules were chosen from 10, 15, 25, or 30 seconds and were constant for a block of rewards. Multiple schedules allowed us to diversify the response dynamics of the animals^7^. Each experimental session contained 2–4 blocks with 34 or 66 rewards in each block. Given the constant hazard rate and the fact that rewards never disappeared once available, the probability of reward availability increased exponentially toward 1 with the time elapsed since the last press (the *waiting time*), with a time constant given by the reward schedule (see Methods; Fig 1d and S2). Since the monkey chose when to respond, its decisions influenced the probability of reward availability (Fig. 1c). An ideal observer that did not know the schedule or availability should track the time and reward histories, so we hypothesize that animals attempt to maximize their reward by tracking these quantities, referred to as the reward predictors, to determine when and where to respond.

We examined if the firing rates of neurons in dlPFC represent the reward predictors as they are continuously evolving in time (monkey G: 11 sessions; monkey T: 19 sessions, *n* = 1323 single and multi-units). Additionally, we extracted the neurons’ press-locked events, *i.e*., firing rates a few seconds before and after each press (Fig. 1b). The Continuous-time neural activity allowed us to understand how continuous representations of task variables in dlPFC leads to the animal’s choice to press. The press-locked neural activity explained how the state of these representation, prior to a press, combined with the new information, which is the reward outcome, predict where and when the animals respond. The continuous spike raster and the press-locked firing rate of a sample neuron (Fig. 1e) is shown for four consecutive box presses with different reward/choice outcomes: an unrewarded press followed by a switch to the other box, an unrewarded press when the animal stayed at the same box, and two rewarded presses when the animal stayed at the same box. The fourth outcome, switching to the other box after a rewarded press, accounted for only 2% of the presses, so we do not show it in this example.

Before attempting to understand which neural events predicted when and where to press, we attempted to identify variables that the animal can either observe or control and that they potentially use to estimate the chances of rewards. Consequently, we determined if these variables empirically predicted the next reward outcome in our experiment. We also tested whether they predict the animal’s choice of box and time of press. Next, we identified the neural representation of these variables in the population of recorded neurons in dlPFC. Finally, we tested whether these representations predicted the animal’s choices in advance.

### Predictors of the next reward

According to the marginal value theorem of foraging theory^1^, an animal could optimize its reward while minimizing travel costs by estimating the box schedules, tracking the temporal evolution of the probability of reward availability, and using them to choose when and where to search for reward. Although the probability of the reward availability is the best predictor of the randomly generated reward, it was completely unobservable to the animals in our experiment. However, other predictive variables were observable or controllable by the animals, such as the waiting time between the presses or the reward ratio, defined as the proportion of current option’s recently delivered reward from the total of recently delivered rewards. The *recency* criterion was imposed by applying a causal Gaussian filter to the binary sequency of delivered (1) or denied (0) rewards^5,6^. The waiting time, together with the scheduled reward rate, determines the probability of reward availability (see methods; fig. S2). The reward ratio, when tracked on a time-scale relevant to the volatility of the environment^5^, is a proxy for the scheduled reward ratio, defined as the ratio of the scheduled reward rate on the current box and the sum of the scheduled reward rates of two boxes. Because the scheduled reward ratio changes without warning from block to block, we maximized the correlation of the scheduled reward ratio with the animal’s observed reward ratio by tuning the width of the causal Gaussian filter mentioned above (Fig. S3).

We assessed how well each variable predicted the reward by correlating the rewarded fraction of presses with that variable prior to each press. Specifically, we pooled 8862 behavioral presses from 30 sessions of two monkeys, binned them according to each hidden or observable/controllable variable so that there were 50 presses in each bin, calculated the fraction of rewarded presses within each bin (Fig 2a), and computed the Pearson correlation between the binned variable and rewarded fraction of presses. Naturally, the probability of reward availability was highly correlated (*r*=0.93; Fig 2a) with the rewarded fraction of presses. The scheduled reward rate was correlated with the fraction of rewarded presses as well (*r* = 0.43; Fig 2a). This correlation is weaker than the correlation of the waiting time with the fraction of rewarded presses (r = 0.92; Fig 2a) because the probability of reward availability is determined by both waiting time and the scheduled reward rate, and the animals choose a wide range of the waiting times, diluting the prediction of the scheduled reward rate alone.

**Figure 2.**
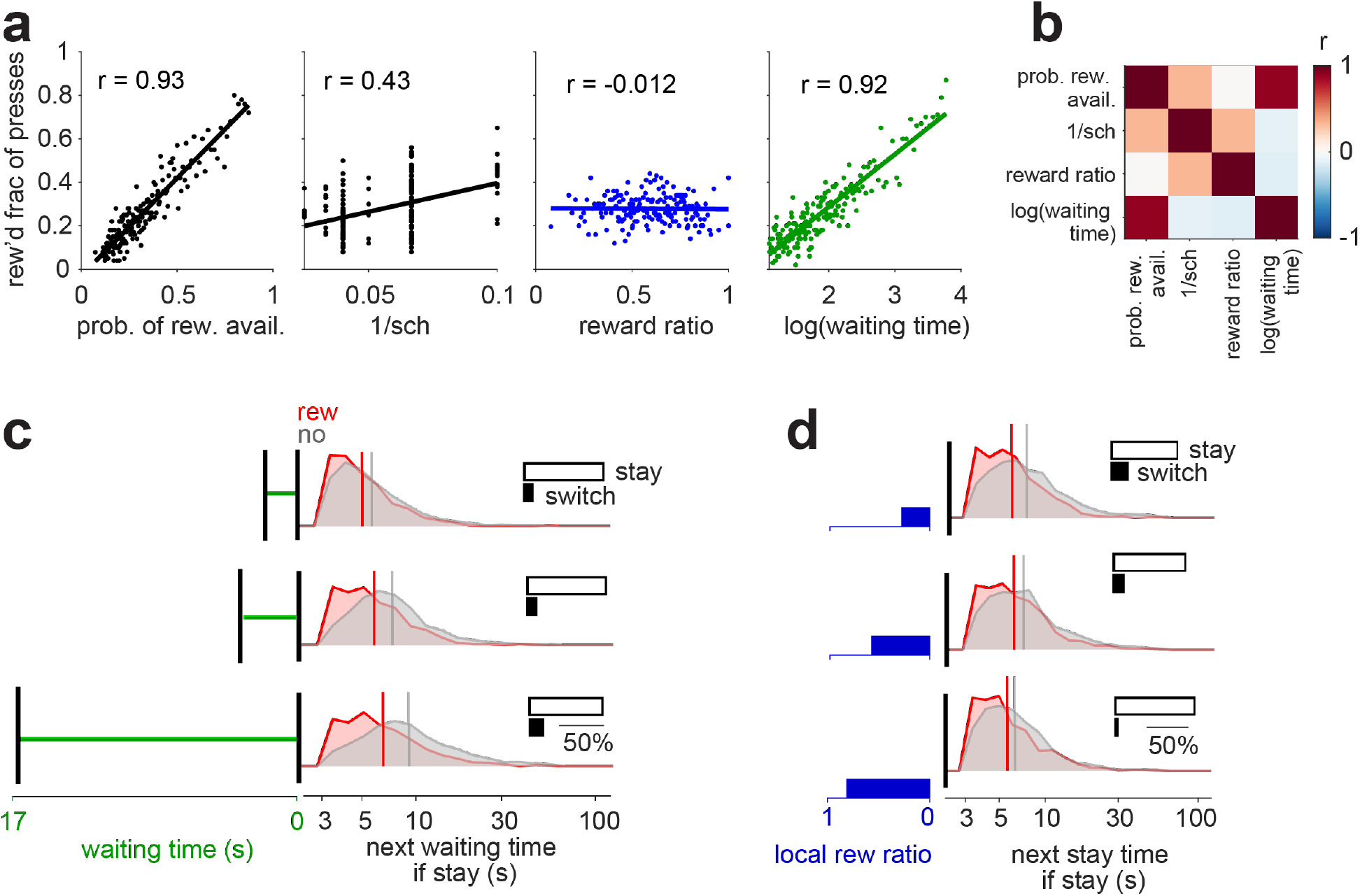
Reward predictors, together with the reward outcome, determine the choices and the next waiting time. **a)** predictability of the next reward from experimental and behavioral variables. 8862 presses from 30 sessions were pooled together and binned into 50-press bins according to each experimental variable. The rewarded fraction of presses was calculated in each bin, then the Pearson correlation coefficient was calculated across bins between the average of the experimental variable and the rewarded fraction of presses. **b)** correlation matrix of the task variables in a. **c)** The histograms of the next waiting time for rewarded and unrewarded presses that were made after a short, medium, or long wait, determined by equal intervals in the percentile of the presses. *Inset*: Increase in probability of switching, when not rewarded after a short, medium, or long wait. The probability of switching after being rewarded was less than 2% and therefore excluded from this analysis. **d)** same as c but for reward ratio instead of waiting time.

Although the waiting time was highly predictive of the next reward, the reward ratio potentially plays an important role in animal’s subjective reward expectation^6^. The reward ratio was not correlated with the fraction of rewarded presses (Fig. 2a, r = −0.012). However, it was positively correlated with the scheduled reward rate on the side that the animal pressed (*r* = 0.32), meaning that it might be used by the animals as an observable estimation of the hidden reward rates. Moreover, it was only weakly correlated with the log of waiting time (*r* = −0.14), meaning that it may be considered by the animals as a source of information, independent from the waiting time. We refer to the waiting time and the reward ratio as the reward predictors because they may be used by the animals to predict the reward, therefore may play a role in determining the animals’ reward expectation (See Fig. S4 for the analysis of other observable reward predictors).

### How do reward predictors determine ‘when’ and ‘where’ to press?

Although the subjective reward expectation is not directly measurable, we might infer changes in the animals’ reward expectation from the animals’ next choice, after reward is delivered or denied. For example, an animal may realize that waiting longer increases its chances of receiving a reward, so we expect that an unrewarded press after a long wait might lead it to wait even longer between presses at the current box. Alternatively, the animal may realize that the waiting time for getting reward at the current box is too long. Therefore, it may switch to the other box anticipating a better reward rate. We thus hypothesized that the animals’ decision on where and when to press depends upon the reward predictors, as the bases of animals’ reward expectation. We evaluated this hypothesis by analyzing the effect of such reward predictors on the probability distribution of the next waiting time and the probability of switching. These events were grouped depending on whether presses were rewarded and occurred after a short (3-5 s), medium (5-8 s), or long (8-60 s) wait (Fig. 2c). An unrewarded press increased the next wait by 10% (AUC = 0.53 ± 0.03) after a short wait, by 28% (AUC = 0.59 ± 0.02) after a medium wait, and by 42% (AUC = 0.59 ± 0.02) after a long wait, each compared to the corresponding average waiting times for rewarded presses. Moreover, the probability of switching to the other box increased with the duration of unrewarded waits (9.5%, 10.2%, and 16.5% more switches after a short, medium, and long waiting time; Fig. 2c, *insets*). These choice differences (to continue pressing the button for the same box or switch to the other box) and the next waiting time when choosing to press on the same box, demonstrate that animals base their expectation of reward on their waiting time and adjust their behavior by waiting longer before the next press or switching to the other box when this expectation is not met. While previous studies point to melioration, that is following the current flow of reward delivery^4^, we provide evidence of more temporally structured computations: the animals predict the chance of the next reward as they choose how long to wait before making the next press and adjust the waiting time when their expectation is not met. A key to this finding was a trial-free task, allowing animals to experience a wide range of waiting times and discovering that longer intervals yielded a higher chance of receiving a reward.

The animals might also develop expectations about the quality of the current box from the reward ratio. Again, we can infer these expectations indirectly through changes in the next waiting time and choices. After unrewarded presses, animals waited longer and switched more, with the smallest changes for biggest reward ratios (Fig 2d; 23%, 18% and 15% longer unrewarded waits and 19%, 12% and 3% switches after a low, medium, and high reward ratio). This suggests that animals require stronger evidence to override a better reward history.

Altogether, this provides evidence that an animal’s policy on when and where to press depends on whether the box delivers a reward, as expected after a long waiting time or a high reward ratio. We inferred that animals update their expectation when those expectations are violated by the lack of an expected reward. This policy is a case of “learning a guess from a guess”^25^ which is useful in the absence of sensory evidence directly cueing the probability or availability of reward. To provide further evidence that the waiting time and reward ratio underlie animals’ reward expectation, we examined their encoding in the recorded neural population.

### Decoding reward predictors from the neural population

Prior to a motor action, the activity of neurons in dorsolateral prefrontal cortex (dlPFC) is correlated with the value of a visually cued expected reward^16^ or the probability of reward, estimated by the recent history of reward delivery^10^. Therefore, we hypothesized that the activity of dlPFC neurons, prior to each press, encodes the reward expectation for that press, for the range of the reward predictors variables observed or generated by each animal. For example, the neuron in Fig. 3a *left* responds more before a press following a long wait (top 20% of waiting times in that session) compared to a short waiting time (bottom 20%; Wilcoxon rank-sum test, *p* ≪ 10^-3^). Similarly, the neuron in Fig. 3a *right* responds more when the reward ratio prior to a press is in the bottom 20% compared to when it was in the top 20% (Wilcoxon rank-sum test, *p* ≪ 10^-3^).

**Figure 3.**
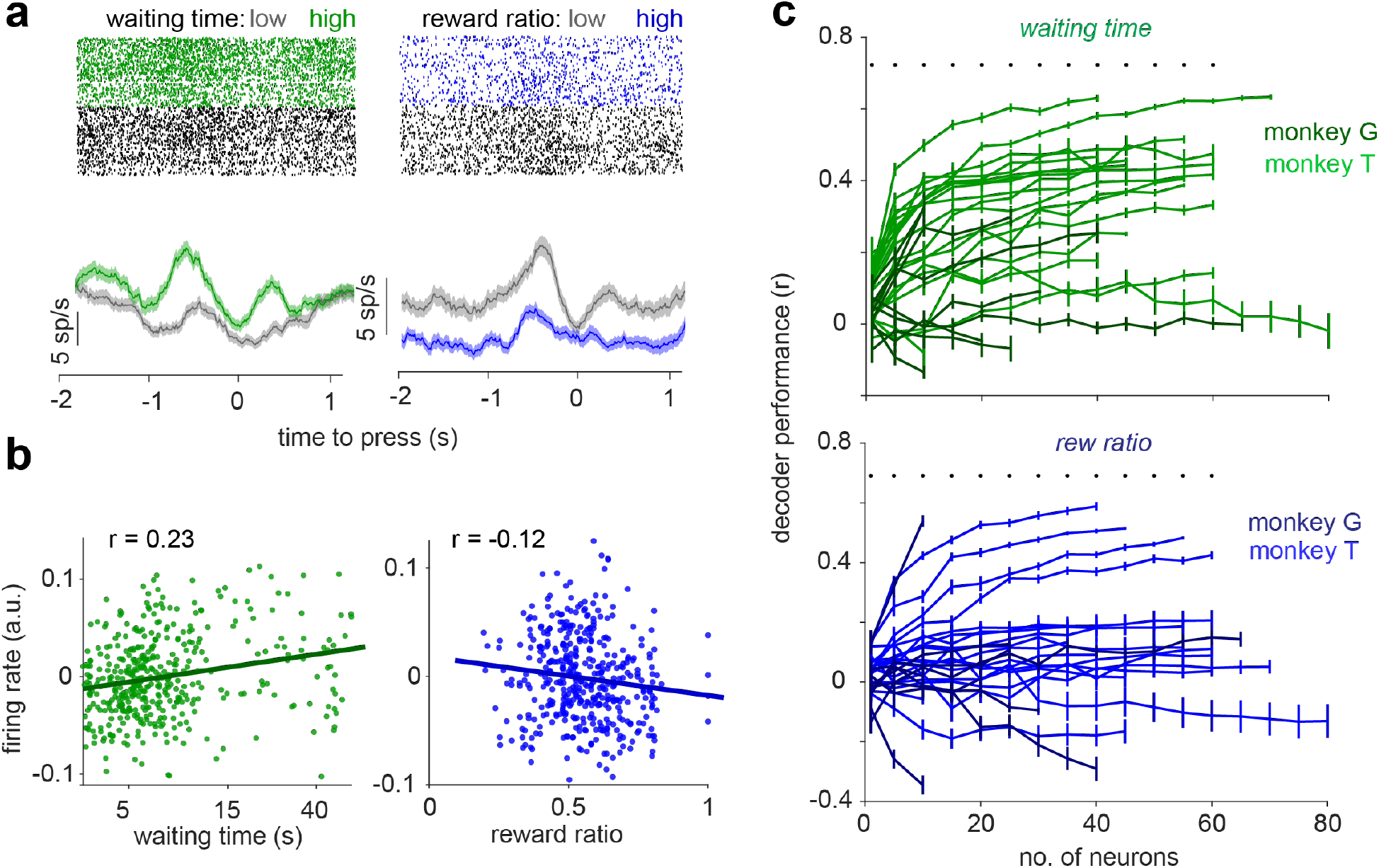
Neuronal populations encode variables of the reward dynamics. **(a)** Sample neurons for which the pre-press firing rate covaries with either waiting time (*left*) or reward ratio (*right*). The firing rate was calculated for a 200 ms sliding windows starting 2 s before and ending 1 s after presses. Firing rates were averaged over presses with low (< 20^th^ percentile, gray) and high (> 80^th^ percentile, colored) of either the waiting time or the reward rate. **(b)** For the neuron in a, the scatter of firing rates, after decorrelating them from the locomotion, as a function of the waiting time (*left*) or the reward ratio (*right*). **(c)** Prediction of the waiting time (*top*) or the reward ratio (*bottom*) for 30 sessions as a function of the number of neurons used as predictors. The predictor neurons were chosen randomly from the population. The random selection was done 20 times for each data point. Sessions of monkey G are shown with a darker color and sessions of monkey T are shown with a lighter color.

Because task-irrelevant variables such as locomotion, limb and eye movement and pupil size before or after presses may influence dlPFC activity, we performed control experiments to quantify the correlation between task-irrelevant variables and neural activity. First, our control experiments in which animals moved to receive reward from the same boxes as in Fig. 1a revealed that eye movements have only a minor influence on neuronal activity while animals interacted with the box, although they have a stronger influence during locomotion (*r* = 0.16, *p ≪* 10^-3^, for eye velocity, and *r* = 0.13, *p ≪* 10^-3^, for fixation rate ^23^). We thus decorrelated the neural activity from the locomotion by projecting neural activity onto the subspace orthogonal to locomotion (see Methods) such that the remaining neural activity was uncorrelated with locomotion (Fig. S5). Second, one animal performed the same task as presented here, while its arm movements, pupil diameter, and eye velocity were monitored using the same eye tracking method as in^23^. We found ≤ 9% of neurons in dlPFC with significant (p<0.01) correlation with the arm movement (Fig. S6) in 1 s time intervals starting 2 s before and ending 2 s after presses. Pupil diameter was correlated with ≤ 10% of neurons. However, after we decorrelated the neural activity from the locomotion, the percentage of neurons with a significant correlation with the pupil diameter dropped to ≤ 7%. Similarly, the percentage of neurons with a significant correlation with the eye velocity dropped from ≤ 9% to ≤ 4%. Because decorrelating the neural activity from locomotion also decreases the correlation between the neural activity and other task-irrelevant variables, we focused our analysis for the rest of this study on the neural activity that was decorrelated from the locomotion.

We further examined whether the neural activity before a button press encodes the wait time preceding that press. Modulation of pre-motor cortical activity with the value of a motor action, here determined as the chance of receiving a reward, can improve action execution for rewarding actions. Therefore, finding a representation of an entire spectrum of waiting times may be significant as it provides evidence that animals assign a value to a button press proportional to the theoretical chance of reward delivery which monotonically grows with waiting time. We first measured the spike counts in a 1-second interval (*i.e*., a “pre-press” interval from –1.1 to –0.1 seconds) for each neuron (*n* = 1323 single and multi-units). This time interval was selected since the arm movement starts approximately 0.5 s before the press is recorded, and the modulation of neural activity typically starts around 0.5 s before that movement^26^.The pre-press firing rate of the neuron in Fig. 3a *left* was correlated with the waiting time (Spearman correlation coefficient = 0.24; p≪ 10^-3^; Fig. 3b *left*). For the entire population of cells, around 35% of neurons exhibited a significant Spearman correlation (*t*-test, p < 0.01; 31% positively correlated and 4% negatively correlated; Monkey G: 27%; Monkey T: 37%).

To further examine how information about the waiting time is distributed across neurons, we decoded the waiting time from population activity prior to each press using the spike counts of randomly subsampled sets of neurons (see Methods for a description of the regression-based decoder analysis). Our decoder analysis revealed that even random neural subpopulations encode the waiting time (Fig. 3c; p<=0.01).

Furthermore, consistent with previous reports^6,10,17,27^, we found that dlPFC neurons encode the reward ratio. Over the entire population, there was a significant correlation between the pre-press firing rate and reward ratio (*t*-test, *p* < 0.01) for 23% of the neurons (9% positively correlated and 14% negatively correlated; Monkey G: 12%, Monkey T: 26%). Decoder performance for the reward ratio was higher than chance (p<0.01) when we used a sub-population of 1 or more neurons as the predictors. Taken together, these results indicate that both reward predictors are encoded in the pre-press neural activity at the individual neuron and population levels. This finding provides further evidence that the animals’ reward expectation is founded on the chosen reward predictors.

### Identifying continuous representation of task variables in a latent space

A notable difference between the waiting time and reward ratio is that the reward ratio jumps discretely at press times while the waiting time, by nature, evolves continuously. To measure the degree to which the waiting time explains the variability in the continuously evolving activity of neurons, we attempted to fit the variability in a simultaneously recorded population of neurons at any given point in time using a weighted sum of task-relevant variables, passed through a set of temporal basis functions^28–30^. For event-based task variables such as presses, rewards, and switches (pressing the button on the other box), each event raster was convolved with temporal basis-functions (Fig. 3e). We used pulse shaped temporal basis-functions, each with pulse width of 200 ms. Although simple, these basis functions capture the variability of neural activity effectively. We used different numbers of these pulse basis functions for different variables: seven basis functions for the pre-press time-interval, post-press time, and post-choice; and ten basis functions for the post-reward time (spanning 2 s; Fig. 4a). For continuously evolving task-variables — the waiting time and the location — we used monomial basis-functions with powers of ½, 1, 2, 3 and 5 (Fig. 4a). The reward ratio was also treated as a continuous task variable for which the value only updates at each press. Neural activity was smoothed by 1 s sliding windows. Locomotion-related neural activity was subtracted as described above. To concentrate our analysis on times when animals were engaged in the task, we excluded time bins preceding or following any presses by more than 5 s.

**Figure 4.**
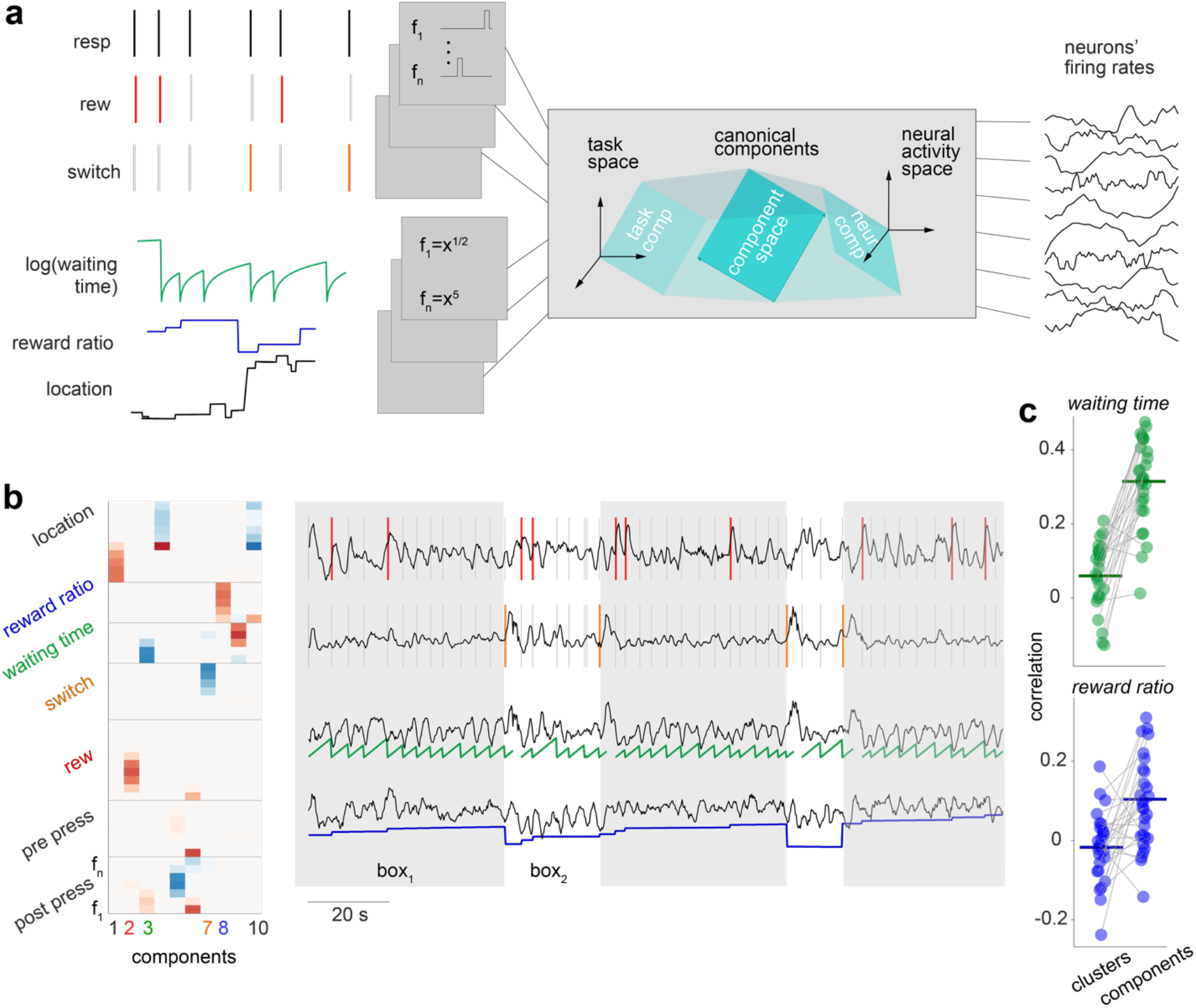
Canonical components of the neural population represent task variables in continuous time. **(a)** Illustration of the canonical correlation analysis for finding a reduced-dimensional space in the task space and the corresponding subspace in the neural activity space. The canonical components define the maximally correlated subspaces between the task variables and the neural activity. The 51-dimensional task space was made from 6 task variables, by passing each variable through a set of basis functions. The basis functions were temporal pulse-shaped filters for press, reward, and choice events (the time of the reward and the choice events were assumed to overlap the time of the press after which the reward was delivered, or the choice was made). The basis functions for continuously evolving task variables (the waiting time, the reward ratio and 2-dimensional location) were a set of power functions with powers of ½, 1, 2, 3 and 5. Overall, 51 predictors were made using these 6 task-variables. The neural space was made using all simultaneously recorded neurons. **(b)** *Left*: The weight of the contribution of each task variable in 10-first canonical components, sorted in the descending order of the correlation between the projection of each component in the task and neural spaces. The indices of the components representing waiting time, reward ratio, reward and choice are color coded for easier association. *Right*: neural representation of four task variables, reward, choice, waiting time, and reward ratio for the same sample session on the *left*. The component that was associated with each of these four task variables was identified as the component for which the absolute value of the weights was highest, compared to the weights for the other task variables. **(c)** Cross-validated Pearson correlation coefficient between the reward predictors, waiting time (*top*) and reward ratio (*bottom*), with either the clusters of 5 or more neurons (Fig. S7) or the canonical components.

To identify the latent representation of task variables in the neural space, we used canonical correlation analysis (CCA) to find components that are shared between the task and the neural spaces. CCA finds these canonical components by applying singular value decomposition to the cross-correlation matrix between two spaces^31^. To favor interpretable latent components such that each component is associated with a small subset of the task variables, we imposed a sparsification penalty (LASSO with fullness constant = 0.3 on the weights of the task variables^31^. This regularization should also help reduce over-fitting of the model. We calculated 10 components for each training set (Fig. 4b), and then identified neural components making the greatest contributions to rewards, choices, waiting time, and reward ratio. Interestingly, the waiting time neural component ramps up between the consecutive presses (Fig. 4c, 3^rd^ row), suggesting that the latent representation of the waiting time might be used by the brain to generate the next press, in a similar fashion to the evidence accumulation models proposed in decision-making ^32^. The reward ratio component followed the difference between the reward ratio of the boxes (Fig. 4c, 4^th^ row). The reward and choice components showed sharp post-press elevated activity (Fig. 4c, 1^st^ and 2^nd^ rows).

We asked whether fitting a model to reconstruct the activity of individually recorded neurons^29^ or sites^28^, then clustering the neurons based on the similarity between the reconstructed activity (Fig. S7) yields a better representation than the latent variables that we found using the canonical correlation analysis. To this end, we calculated the Pearson correlation coefficient between reward predictors and their associated canonical components analysis and compared them with the correlation between the reward predictors with the neuronal clusters in each session that was maximally correlated with each reward predictor. The average correlation coefficient between either of the reward predictors and the neural components was higher than that with the neural clusters of > 5 neurons and the same reward predictor (p ≪ 10^-3^ for the waiting time, p=0.002 for the reward ratio; Fig 4c). This indicates that the latent neural components provide better correlates of reward predictors relative to average activity of groups of neurons that were clustered together based on their task-relevant activity.

### Predicting reward, choice, and the next waiting time

Since the animal cannot know the true latent reward dynamics, its choices can only be driven by its subjective beliefs about these variables, rather than the objective truth from the experiment. For example, imagine that at some time the monkey overestimates the probability of reward availability, whether due to over-estimating the time that he has been waiting or due to over-estimating the scheduled reward rate. He might then be more likely to switch boxes after an unrewarded push than if he had known the true (lower) value of this probability, or than if he had underestimated it. Therefore, we predicted the animal’s choice to switch sides based on neuronal components corresponding to the task variables: we interpret those neuronal components as current estimates of the animal’s subjective beliefs. We decoded the pre-press neural activity by projecting the population activity onto the subspace formed by the first ten canonical components for the reward predictors. This projection accounts for latent representation of reward predictors that could potentially influence the choices or the next waiting time or predict the eventual reward outcome.

We attempted to predict rewards, choices, and the next waiting times from three distinct types of predictors: (i) the pre-press reward predictors (canonical components in the reward predictors’ space), (ii) neural representations of the reward predictors (canonical components in the neural space), and (iii) the entire simultaneously recorded neural population (Fig. 5a). For a fair comparison between the components and the entire neural population, we did not sparsify the weights of task variables in canonical components. To predict the reward, we trained binomial logistic regressions on the same data used to find the canonical components, then tested on the held-out data. To assess the prediction performance, we calculated the area under the R.O.C. curve showing the discriminability of the predictors’ output for the rewarded presses from the unrewarded presses. The same method was used for the choice to stay or switch. To predict the next waiting time, we used generalized linear models instead of the logistic regression and evaluated the performance by calculating the Pearson correlation coefficient between the real and the predicted values. All predictors were trained and tested for each 200 ms time bin, starting 3 s prior to each press and ending 1 s after.

**Figure 5.**
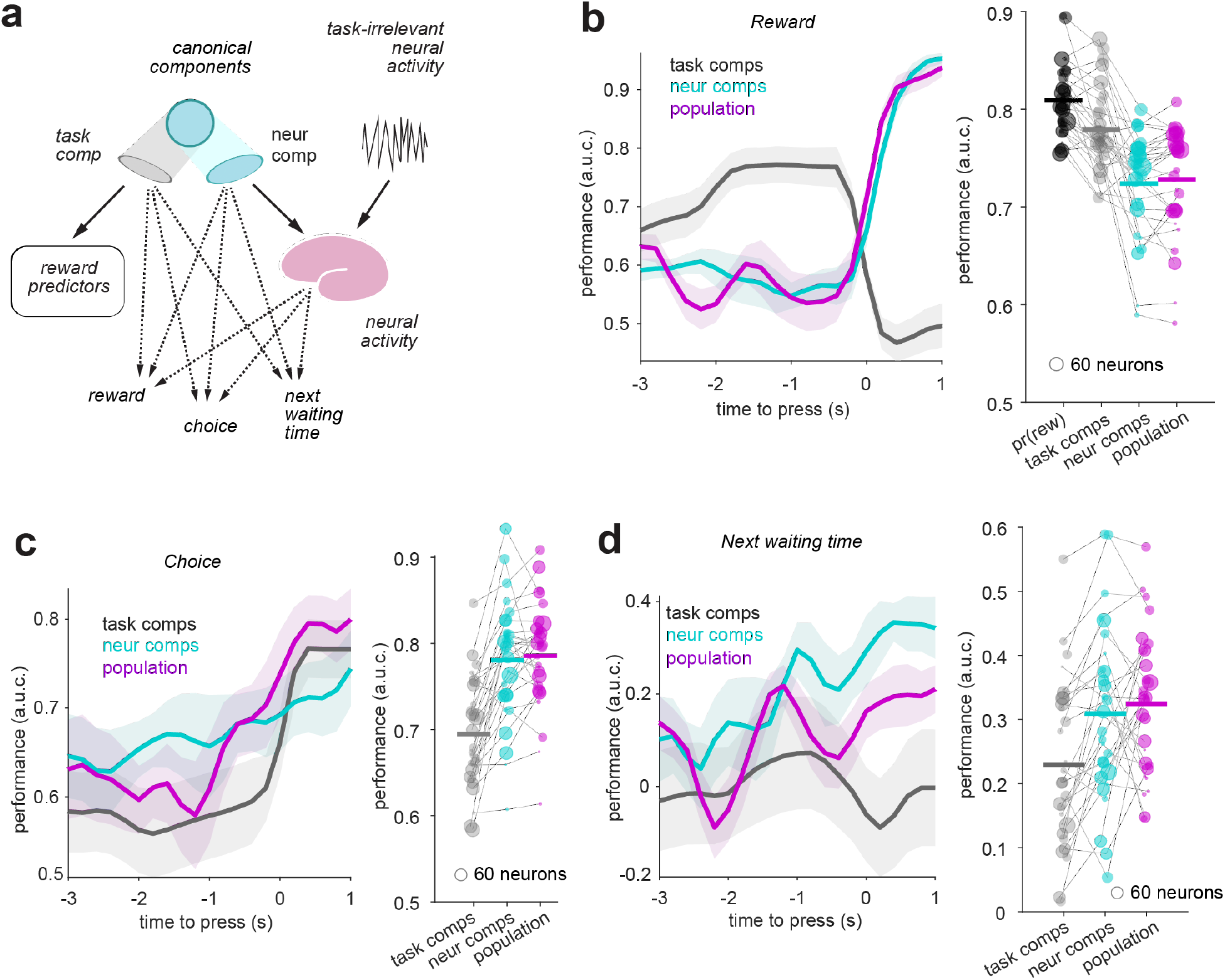
(a) Prediction of rewards, choice, or next waiting time from: (i) task components, (ii) neural components, and (iii) the entire simultaneously recorded population. The task components are defined as the projection of the canonical component in the 10-dimensional space of waiting time and reward rate (each passed through 5 basis functions), The neural components are defined as the projection of the canonical component into the neural population space. The prediction was trained and tested for each 200 ms time-bin, starting 3 s before and ending 1 second after the presses. **(b-d)** The prediction results shown for example sessions (*left*) or summarized across all sessions for the peak within the 2 s time-interval prior to the presses (*right*). *Left*: The prediction performances using the post-press components or population activity are shown for comparison. *Right*: The peak was calculated as the average of five time-bins with the highest prediction performances. The prediction performance for the pre-press time-bins provide evidence that the post-press choices or the rewards were in fact predictable, even before the presses were made.

In the example session shown in Fig. 5b *left*, the prediction of the reward outcomes using the task components improved as the analysis windows approached the time of the press. The reward outcomes are determined by the actual experimental task variables, and indeed we confirmed that the true pre-press task components, the projection of the canonical component in the space of reward predictors, predict reward better than their neural representations, the projection of the canonical components on the neural population space, or the entire neural population (Fig. 5b *right*).

In contrast, the choices and the next waiting time should follow the animal’s subjective estimation of the reward dynamic variables. Fascinatingly, and somewhat unexpectedly, the neural activity before a press predicted the subsequent choice (Fig. 5c) and waiting time (Fig. 5d). Because the animal’s movement to switch to the other box or press the button again occurred after the current press (Fig. S8), the prediction of either of these actions by the neural components precedes the execution of the predicted actions. Therefore, we provide further evidence that the animals construct an expectation of reward prior to a press, based on their subjective understanding of the temporal structure of the task. Subsequently, animals decide when and where to press next based on the expected reward and the actual observed reward. Interestingly, the 10-dimentional neural representation of the pre-press task components predicted the choice and the next waiting time as well as the entire neural population (Fig. 5b-d), indicating that these few canonical neural components successfully capture the relevant signals within the larger neural population dynamics.

It might seem obvious that neural features should be better predictors of when and where to press than experimental variables: after all, the animal’s brain is making its choice and not the experimental equipment. However, it is not evident *a priori* that the relevant neural representations would be found within our recorded dlPFC population, nor whether we record enough neurons to capture enough of the animal’s choice-relevant information. And even if dlPFC does contain the choice-relevant signals, it is not obvious that the neural components for our specific hypothesized reward predictors would be the right ones to predict the choices. It is thus noteworthy that these neurally decoded reward predictors predict choices significantly better than the task variables from which they are derived, and as well as the full neural population. Evidently, our analysis identifies a neural subspace containing correlates of latent variables that are relevant for subsequent choices. This subspace also tends to avoid neural dimensions that contain choice-irrelevant variability, since if present these variations would contribute to overfitting and would only hinder our ability to predict choice. We conclude that we are capturing neural correlates of the animals’ subjective beliefs about the latent reward dynamics that inform their choices.

## DISCUSSION

We used a trial-free, unrestrained-animal approach to demonstrate that freely moving monkeys base their foraging strategy on an internal prediction of reward. This prediction is not based solely on the recent history of reward but relies on an internal estimation of the time they have been waiting since the last time they made a choice which determines the probability of reward availability. Indeed, we found that neural populations in prefrontal cortex contain information about reward predictors. Complementary to previous research in restrained animals^6,10^, we revealed that neural signals not only encode reward information, but also significantly predict animal’s choices after each press during foraging. These findings challenge and extend long-standing theories of reward-seeking behavior ^4^ that suggest that animals follow the choice with the maximum recent rate of reward, without constructing a reward model to predict future behavior, according to the matching law^4^.

We argue that matching, while ubiquitous, does not entail a single computational strategy. For our foraging task, matching behavior is consistent with substantially different strategies. One strategy can be simulated using an agent that switches to the other box after the number of unrewarded presses exceeds a noisy threshold (Fig. S9). This strategy corresponds to a basic ‘win-stay / lose-switch’ rule. We implemented this strategy by sampling the threshold from a Gaussian distribution with the same mean and variance as the *loss count* distribution at times when the animal switches sides. Although this agent is blind to both the average reward rate and the probability of the next reward, it still follows the generalized matching law (Fig. S9). The slight under-matching that we observed resembles the behavior of various species in previous studies^4,5^.

We examined a more complex strategy that tracks reward probability and uses foraging theory to make choices by involving three variables: (i) the time since the preceding press and (ii) the variable-interval schedule — which together determine the probability of reward — and (iii) the relative cost of switching locations, which affects the threshold for when to switch. We simulated an agent that follows such a strategy by making choices based on the correct probability of reward availability on both boxes. The agent switches to the other side when the probability of reward availability on the other box exceeds that of the current box by a fixed switching cost, and otherwise waits for the probability of reward availability to increase everywhere (Fig. S9), in accordance with the Marginal Value Theorem of foraging theory^1^. Unlike the first agent, this agent has complete information about the task. Nonetheless, we again observed nearly matching behavior, now with slight over-matching (Fig. S9). These two simulations show that generalized matching law may arise when following a strategy that is either blind to timing, or fully informed. This implies that matching behavior is not, by itself, informative about the underlying strategy or animals’ ability to grasp the hidden rule of the task.

Surprisingly, we found that the targeted representation within the high-dimensional space of neural population activity predicts choice better than the behavioral dynamics and as well as the entire population of recorded neural activity. This is an important confirmation of how targeted dimensionality reduction can reveal neural computations better than behavior or unprocessed neural activity. This type of analysis is essential in natural experiments where task variables are correlated.

One limitation of our findings is the extent to which our results can be generalized across other types of reward dynamics. The reward dynamics in our task are stochastic and time-based, and they resemble the repletion of food resources found in nature. Follow-up studies are needed to determine whether our findings apply to other reward schemes, such as non-Markovian, more clock-like dynamics, or those based on press rate ^33^ whereby reward becomes available after a variable number of presses rather than a variable timeinterval.

Additionally, our study does not determine if the enhanced modulation of neural activity prior to a press is due to a representation of a higher reward expectation or a vigorous motor action due to a higher reward expectation. Comparing the neural modulation between simultaneously recorded populations in dlPFC and pre-motor cortex while continuously measuring the vigor of each press can dissect these two representations.

Finally, by allowing animals to move freely during foraging, our study represents a pioneering move toward studying neural correlates of natural cognition in a free-roaming setting. This paradigm shift has been suggested decades ago^20^, but is only feasible now due to advances in low-power, high-throughput electrophysiological devices as well as large-scale computing^34^. Previous studies have underestimated the cognitive capacity of monkeys during foraging, and this might have primarily been because of their restrictive experimental paradigms. Our freely-moving experimental paradigm likely increases the engagement of natural decision-making processes in the animal’s brain, and reduces the distortions in population dynamics associated with unnatural head-fixed tasks ^35^. The free-roaming setting also enabled us to implement a natural switching cost between two reward options by simply allowing the monkey to walk between them. This is commonly implemented as a timeout period immediately after switching decisions, which potentially alters neural presses in dlPFC. It is also possible that the higher arousal state of the monkey in the free-roaming setting^23^ enhances their cognitive ability to perform the task. Overall, we argue that a shift toward more natural behavior is inevitable for understanding neural mechanisms of cognition^34,36–40^.

## METHODS

All experiments were performed under protocols approved by The University of Texas at Houston Animal Care and Use Committee (AWC) and the Institutional Animal Care and Use Committee (IACUC) for the University of Texas Health Science Center at Houston (UTHealth). Two adult male rhesus monkeys (Macaca mulatta; monkey G: 15 kg, 9 years old; monkey T: 12 kg, 9 years old) were used in the experiments. An additional adult male rhesus monkey (Macaca mulatta; Monkey M, 10 kg, 11 years old) was used for the control experiment, tracking the eye and limb movements.

### Behavioral training and testing

After habituating each monkey in a custom-made experimental cage (120 cm x 60 cm x 90 cm, L×W×H) for at least 4 days per week for over 4 weeks, we trained animals to press the button on each box to receive a reward. Over the course of 4-6 months, we gradually increased the mean time in the VI schedule to let the monkeys grasp the concept of probabilistic reward delivery. Once we started using VI10 (corresponding to an average reward rate of < 0.1 rew/s), monkeys started to spontaneously switch back and forth between the two boxes. If the monkeys disengaged from the task or showed signs of stress, we decreased the VI schedule (increased the reward rate) and kept it constant for one or two days. If the monkey showed a strong bias toward one reward source, we used unbalanced schedules to encourage the monkeys to explore the less preferred box.

After training, we tested monkeys using a range of balanced and unbalanced reward schedules. For balanced schedules we used VI20 or VI30 on both boxes. For unbalanced schedules, we used VI20 versus VI40, VI15 versus VI25, or VI10 versus VI30. The unbalanced schedules may reverse once, twice or three times during a session, e.g. the box with VI20 becomes VI40 and the box with VI40 becomes VI20 after the reversal. Each session lasts until the monkey receives 100 or 200 rewards, ranging from 1-7 hours including a 1-hour break after 100 rewards in sessions with 200 rewards. If monkeys were not engaged with the task for more than 2 minutes, we sometimes interrupted them to encourage them to engage with the task. For the analysis, we exclude all presses which occurred more than 60 s. For the press-locked analysis, we also excluded presses that were made less than 2 s after the previous press to avoid mixing in the press-locked neural activity.

### Analysis of location dependency and locomotion

To determine the physical location and locomotion of the monkey, an overhead wide-angle camera was permanently installed in the experimental cage and the video was recorded at an average rate of 6 frames per second. Each frame was post-processed in six steps using custom-made Matlab code. First, the background image was extracted by averaging all frames in the same experimental session, then it was subtracted from each frame. The background-subtracted image was then passed through a manually determined threshold to identify the dark areas. The same image frame was also processed using standard edge detection algorithms. The thresholded and edge detected images were then multiplied together, and the result was convolved with a spatial filter, which was a circle with the estimated angular diameter of the monkey. The peak of this filtered image was marked as the location of the monkey. We used this heuristic because the illumination of the experimental room and the configuration of object was constant. We expect novel techniques for motion and posture detection using deep neural network^41,42^ to yield similar results. Locomotion (velocity) was calculated as the vector difference between monkey locations in consecutive frames divided by their time difference.

### Determining the reward availability and calculating probability of reward availability

In each time bin of size *dt* = 10 ms, reward became available at a given box if a sample from a Bernoulli distribution was 1. The probability of this event was *dt/VI* where *VI* is the Variable Interval schedule. When the reward became available, it stayed available until collected by the animal. This makes the probability of reward availability a function of the scheduled variable interval as well as the time since the preceding press:

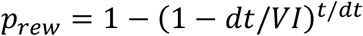

where *t* is the time since the preceding press (Fig S1).

### Chronic implantation of the Utah array

A titanium head post (Christ Instruments) was implanted, followed by a recovery period (> 6 weeks). After acclimatization with the experimental setup, each animal was surgically implanted with a 96-channel Utah array (BlackRock Microsystems) in the dorsolateral prefrontal cortex (dlPFC) (area 46; anterior of the Arcuate sulcus and dorsal of the principal sulcus (Figure S1). The stereotaxic location of dlPFC was determined using MRI images and brain atlases prior to the surgical procedure. The array was implanted using the pneumatic inserter (Blackrock Microsystems). The pedestal was implanted on the caudal skull using either bone cement or bone screws and dental acrylic. Two reference wires were passed through the craniotomy under and above the dura mater. After the implant, the electrical contacts on the pedestal were protected always using a plastic cap except during the experiment. Following array implantation, animals had at least a 2-week recovery period before we recorded from the array.

### Recording and pre-processing of neural activity

To record the activity of neurons while minimizing the interference with the behavioral task, we used a lightweight, battery-powered device (Cereplex-W, Blackrock Microsystems) that communicates wirelessly with a central amplifier and digital processor (Cerebus Neural signal processor, Blackrock Microsystems). First, the monkey was head-fixed, the protective cap of the array’s pedestal was removed, the contacts were cleaned using alcohol and the wireless transmitter was screwed to the pedestal. The neural activity was recorded in the head fixed position for 10 minutes to ensure the quality of the signal before releasing the monkey in the experimental cage. The cage was surrounded by eight antennas. In the recorded signal, spikes were detected online (Cerebus neural signal processor, Blackrock Microsystems) using a manually selected upper threshold on the amplitude of the recorded signal in each channel or an upper and a lower threshold which were ±6.25 times the standard deviation of the raw signal. To minimize the recording noise, we optimized the electrical grounding by keeping the connection of the pedestal to the bone clean and tight. The on-site digitization in the wireless device also showed lower noise than common wired head-stages. The remaining noise from the movements and muscle activities of the monkeys was removed offline using the automatic algorithms in offline sorting (Plexon Inc.). Briefly, this was done by removing the outliers (outlier threshold = 4-5 standard deviations) in a 3-dimensional space that was formed by the first three principal components of the spike waveforms. Then, the principal components were used to sort single units using the expectation-maximization algorithm. Each single and multi-unit signal was evaluated using several criteria: consistent spike waveforms, modulation of activity with 1-sec of the button pushes, and exponentially decaying ISI histogram with no ISI shorter than the refractory period (1 ms). The analyses used all spiking units with consistent waveform shapes (single units) as well as spiking units with mixed waveform shapes but clear pre- or post-press modulation of firing rates (multi-units).

### Removing task-irrelevant components from neural activity

For each neuron *k* we remove movement-related temporal components of the press *rkt*, by subtracting its projection onto the subspace spanned by the task-irrelevant variables: 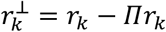, where *⊓* is the projection matrix *⊓* = *L*(*LL*^⊤^)^-1^ *L*^⊤^ and *L* is the *T*×1 vector describing the time series of locomotion, calculated as the magnitude of the changes in the 2-dimensional location.

### Regression-based and binary decoder analysis

To decode a binary variable, such as the reward or the choice to stay or switch, we used logistic regression. To evaluate this model, we used the area under the R.O.C. curve (AUC) to determine the separability of the probability distributions of the held-out samples belonging to either of the classes (reward vs no reward, stay vs. switch). To decode continuous-values variable such as waiting time or the reward ratio, we used a linear regression model^43^. To evaluate this model, we calculated the Pearson correlation coefficient between the measured and predicted values. To train these decoders we divided the presses in each session to 4-18 blocks, holding out one block at a time for testing and using the rest of the blocks for training. To divide the presses into blocks, we found the gaps in press times that were larger than 30 s, then placed all presses between consecutive gaps in one block.

### Canonical correlation analysis (CCA)

Canonical components were calculated using singular value decomposition of the covariance of matrix X in which the columns were task variables, passed through sets of orthogonal basis functions, and matrix Y in which the columns were the pre-press firing rates of simultaneously recorded neurons, for presses within a session. We used custom code in Matlab to call the functions in R ^31^ to calculate canonical components and their correlation coefficients. The cross-validation procedure was the same as for the decoders.

### Statistical analysis

We used the two-sided Wilcoxon signed-rank test except where indicated. We chose this test rather than parametric tests, such as the *t*-test, for its greater statistical power (lower type I and type II errors) when data are not normally distributed. When multiple groups of data were tested, we used the FDR multiple comparisons^44^ correction whose implementation is a standard function in Matlab. No statistical methods were used to predetermine sample sizes. However, the size of our dataset and the number of the experimental sessions greatly exceeded similar studies.

## Contributions

NS, AW, and VD designed the setup and the experiment. NS and AW collected the data. NS, XP and PS performed the analysis. NS, XP, PS, and VD wrote the manuscript. MF collected the supplementary data in Fig. S6. AP and MF analyzed the data for Fig. S6.

## Supplementary figures

**Figure S1.**
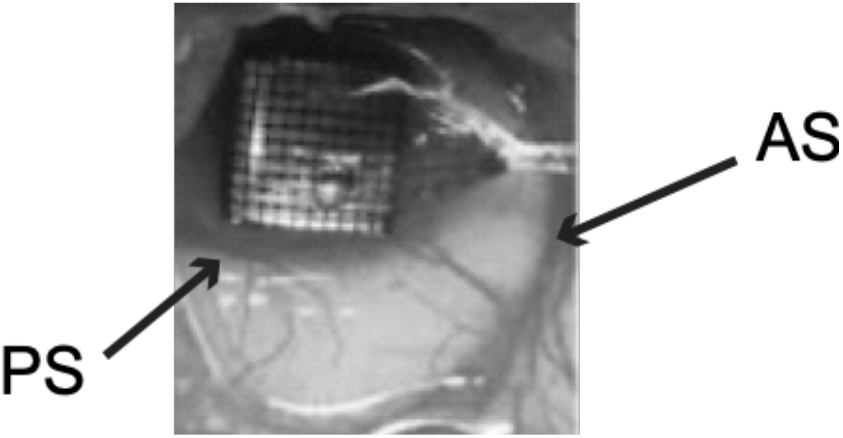
The location of a 96-channel Utah array in dlPFC (area 46) on the left hemisphere of monkey G. The arcuate sulcus (AS) and principal sulcus (PS) are marked.

**Figure S2.**
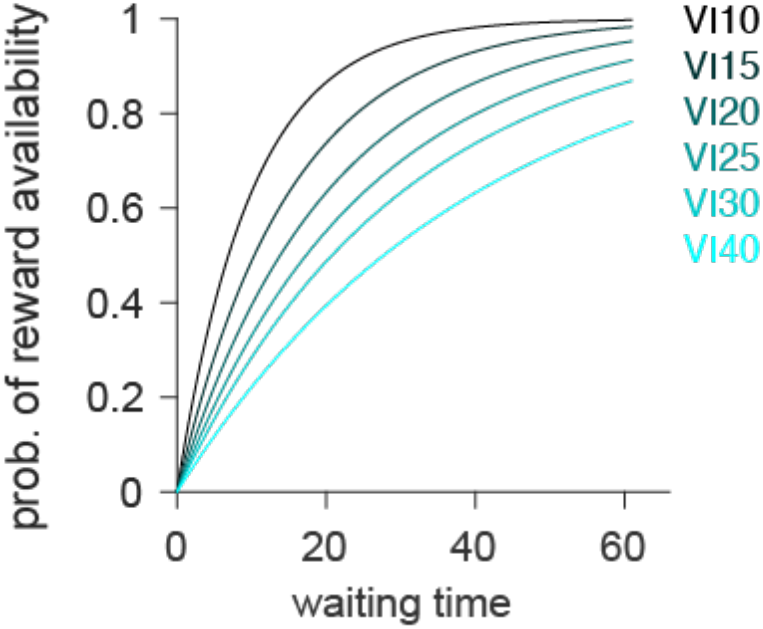
The probability of reward availability as a function of the scheduled reward rate and the time since the preceding response on the same box.

**Figure S3.**
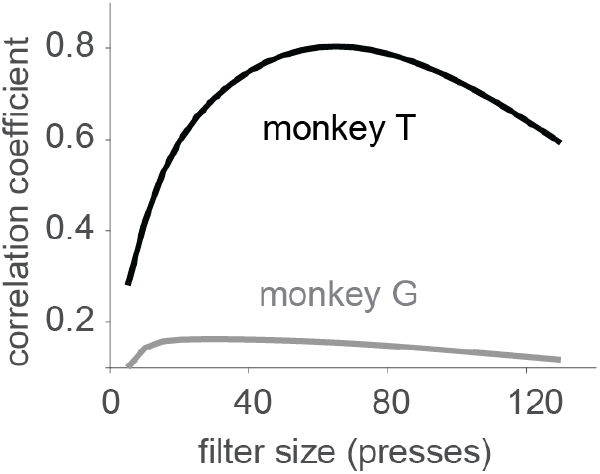
The Pearson correlation coefficient between the scheduled reward ratio and the (observable) reward ratio, calculated using recent sequence of reward outcomes as defined in the main text. The recency was imposed by choosing the standard deviation of a causal Gaussian filter (x-axis). For each monkey, the reward ratio was calculated using the standard deviation for which the maximum correlation with the scheduled reward ratio was achieved.

**Fig S4.**
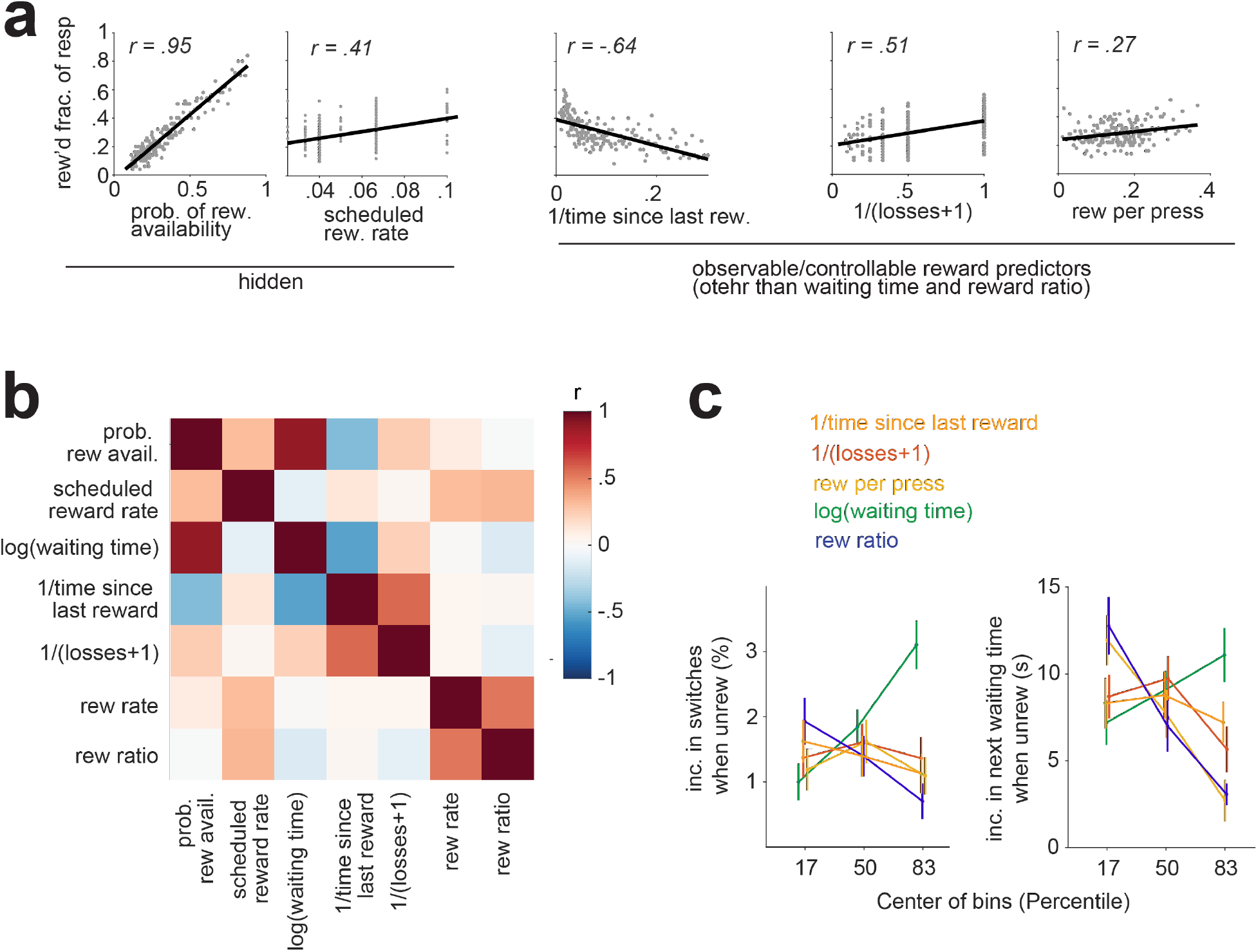
Other observable/controllable variables that might have been used by the animal to predict rewards. **(a)** Similar to Fig 2a, but for the reverse of the time since the last reward, the reverse of the number of un-rewarded presses (losses), and the reward per press (the binary sequency of rewarded (1) and unrewarded (0) presses, filtered using the same causal Gaussian filter that was used for calculation of the reward ratio. **(b)** Correlation matrix between observable and unobservable reward predictors. Based on this matrix, the waiting time was chosen as a reward predictor because it was maximally correlated with the hidden probability of reward availability and the reward ratio was chosen as another reward predictors because it was maximally correlated with the scheduled reward rate and minimally correlated with the waiting time. Other predictors were omitted because they were correlated with either of these two variables (|r|>0.21). **(c)** adjustment on the next action as a function of each of the reward predictor candidates that was discretized into three bins with equal number of presses in each bin. The x-axis shows the center of the bins. The y axis shows the excess percentage of switches or waiting time, when unrewarded, compared to when rewarded. Two chosen reward predictors, waiting time and reward ratio, were linearly predicting the adjustment of the next action.

**Figure S5.**
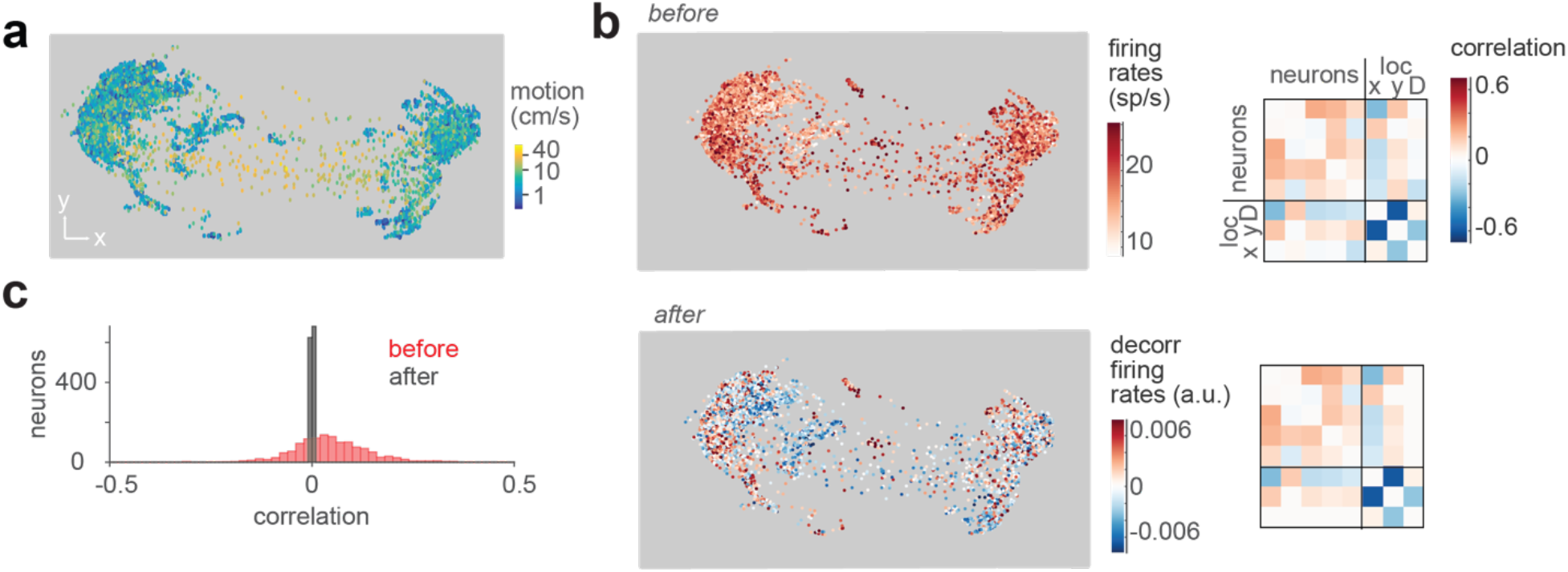
**(a)** Monkey locomotion in a sample session for the press-locked time bins (< 3 s before, or < 1 s after a press). Each dot represents the animal’s position in space, sampled every 200 ms. **(b)** *Left*: Population-averaged firing rates for each time bin in (a) before and after subtracting the vector projection of locomotion. *Right*: Correlation coefficients between locations Loc X and Loc Y, and locomotion Loc D, of the monkey and the pre-press firing rate of each neuron, before and after subtracting the vector projection of locomotion. For a better illustration, an arbitrary subset of neurons is shown. **(c)** Correlation coefficients as in (b) but for all recorded neurons.

**Figure S6.**
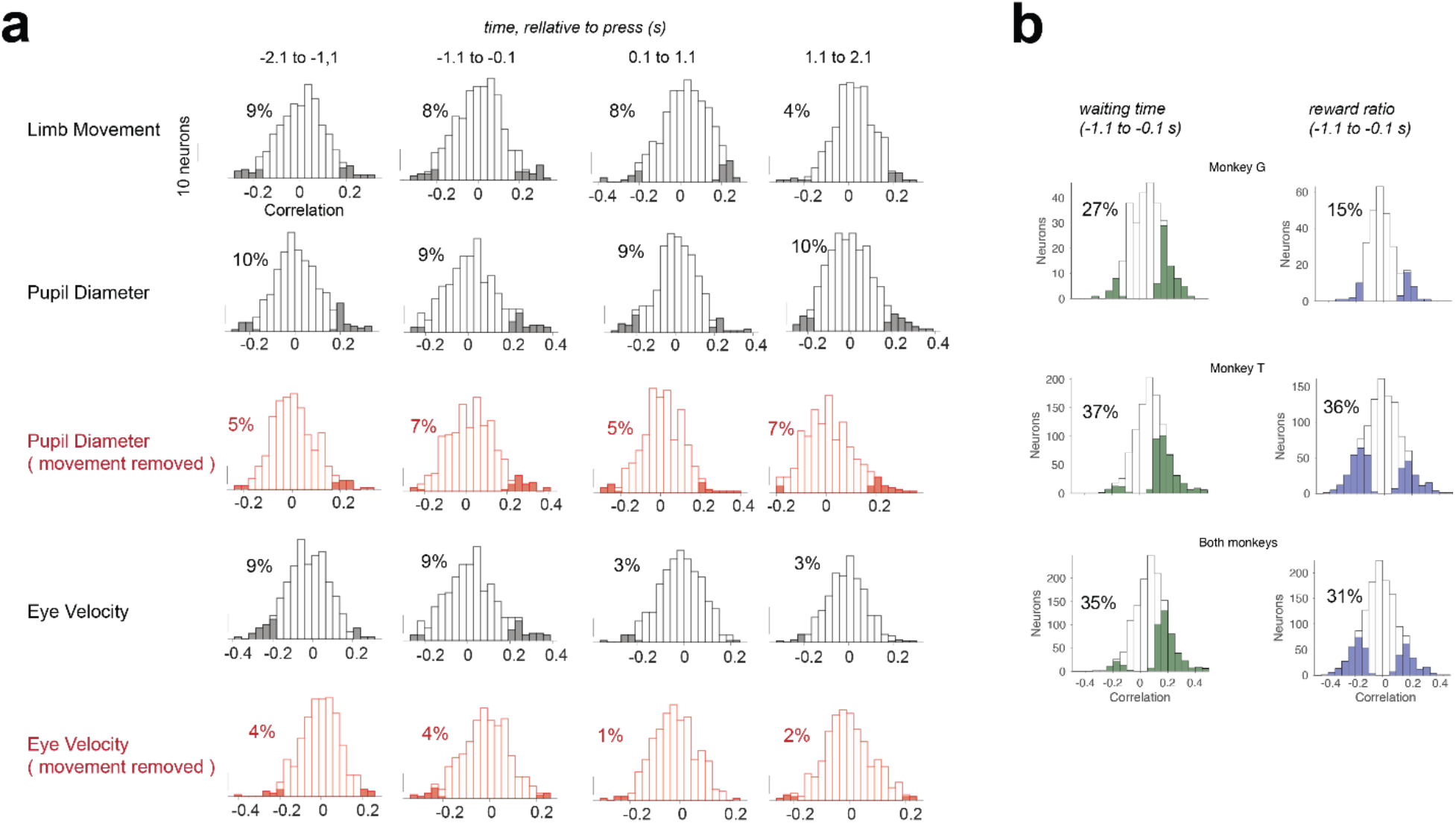
The correlation coefficient between the limb and eye movements and the neural activity of individually recorded neurons. A deep artificial-neural network was trained using DeepLabCut (Mathis et al., 2018) to localize monkey’s shoulder, elbow, and paw of the forelimb contralateral to the recording site (right limb) in each of the frames of the video taken by an overhead camera while the animal performed the foraging task. Using those three markers of the limb in each video frame, we were able to compute average limb movement in any desired time interval. We considered time intervals around button presses (−2s, +2s) and computed the average limb movement and firing rate of all the dlPFC neurons of monkey M in nonoverlapping time-bins of 200-ms width. The pupil diameter and the eye velocity were computed with the same method as^23^. **(b)** For comparison, the correlation coefficient of the reward predictors with firing rates of dlPFC neurons for monkey G and monkey T.

**Figure S7:**
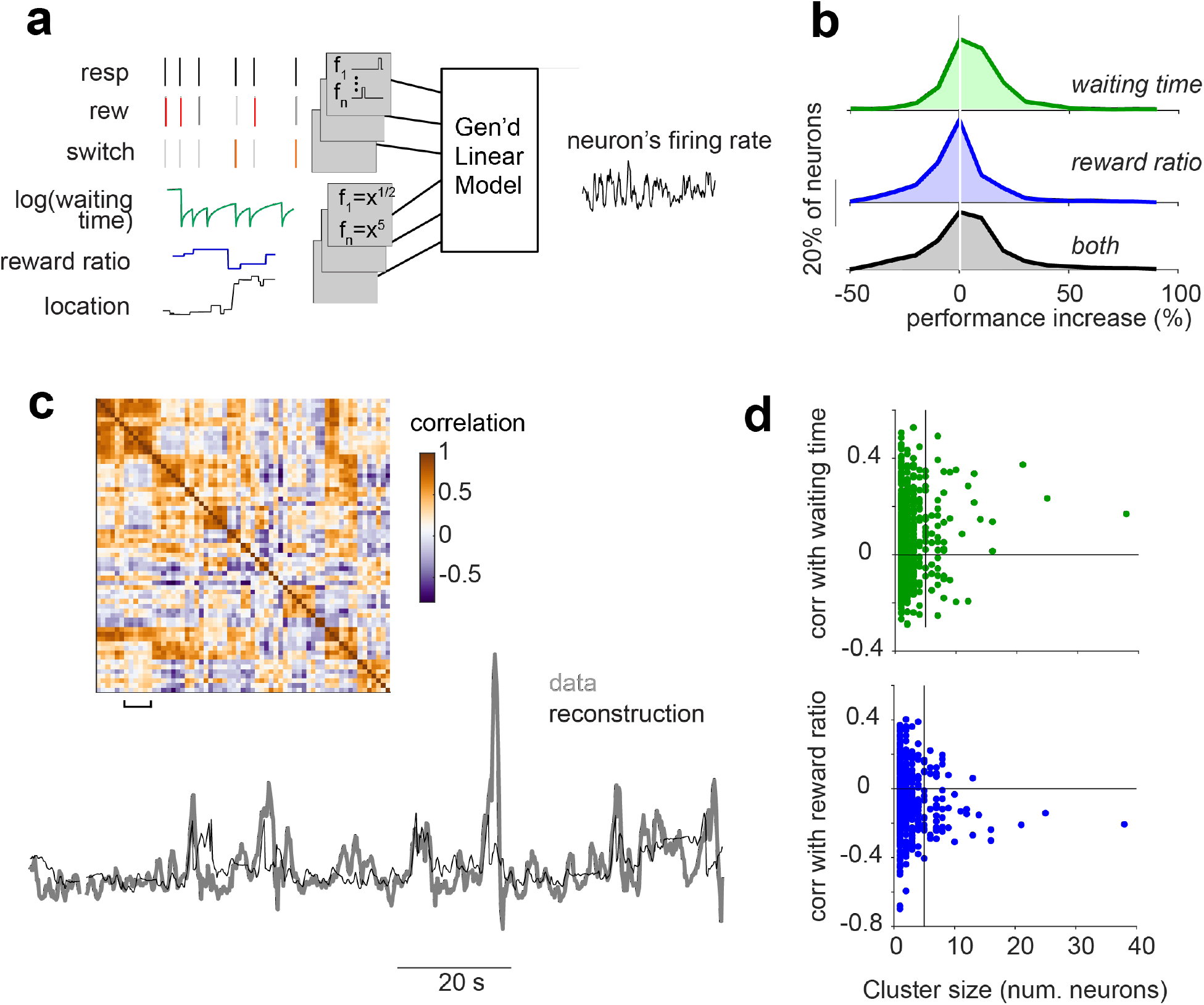
**(a)** A generalized linear model to reconstruct continuously evolving firing rates of individually recorded neurons from the combination of the task variables, passed through a set of basis-functions. The basis functions were pulse-shaped filters for press, reward and choice events (the time of the reward and the choice events were assumed to overlap the time of the press after which the reward was delivered, or the choice was made). The basis functions for continuously evolving task variables (waiting time, reward ratio and 2-dimensional location) were a set of power functions with powers of ½, 1, 2, 3, and 5. Overall, 51 predictors were made using these 6 task-variables. The model was fit to the training data using a Gaussian likelihood function and the trained model was using to reconstruct the neural activity for held-out testing data. **(b)** The improvement in the performance of the model, when either of the reward predictors or both were used, alongside the other task variables in (a), calculated as the percentage increase in the correlation between the recorded and reconstructed activities. While the waiting time improved the model performance for the entire population by 5% (p ≪ 10^-3^), this improvement was insignificant for the reward ratio (p = 1) at the level of individually recorded neurons. **(c)** The reconstructed and recorded firing rates, averaged across 6 neurons in a sample session. The neurons were selected from 60 simultaneously recorded neurons in this session, by clustering the neurons using the correlation matrix of their reconstructed activities, then choosing all neurons in a sample cluster. **(d)** The correlation coefficient between each reward predictor and the reconstructed neural activity that was averaged across neurons in the same cluster vs. the number of neurons in the cluster. The vertical line shows cluster size = 5 which was the criterion that was used to assess the statistical significance of the correlation across the clusters. Across sessions, the average firing rates of clusters of ≥ 5 neurons were positively correlated with the waiting time (p = 0.004, *top*) and negatively correlated with the reward ratio (p = 0.004, *bottom*).

**Fig S8.**
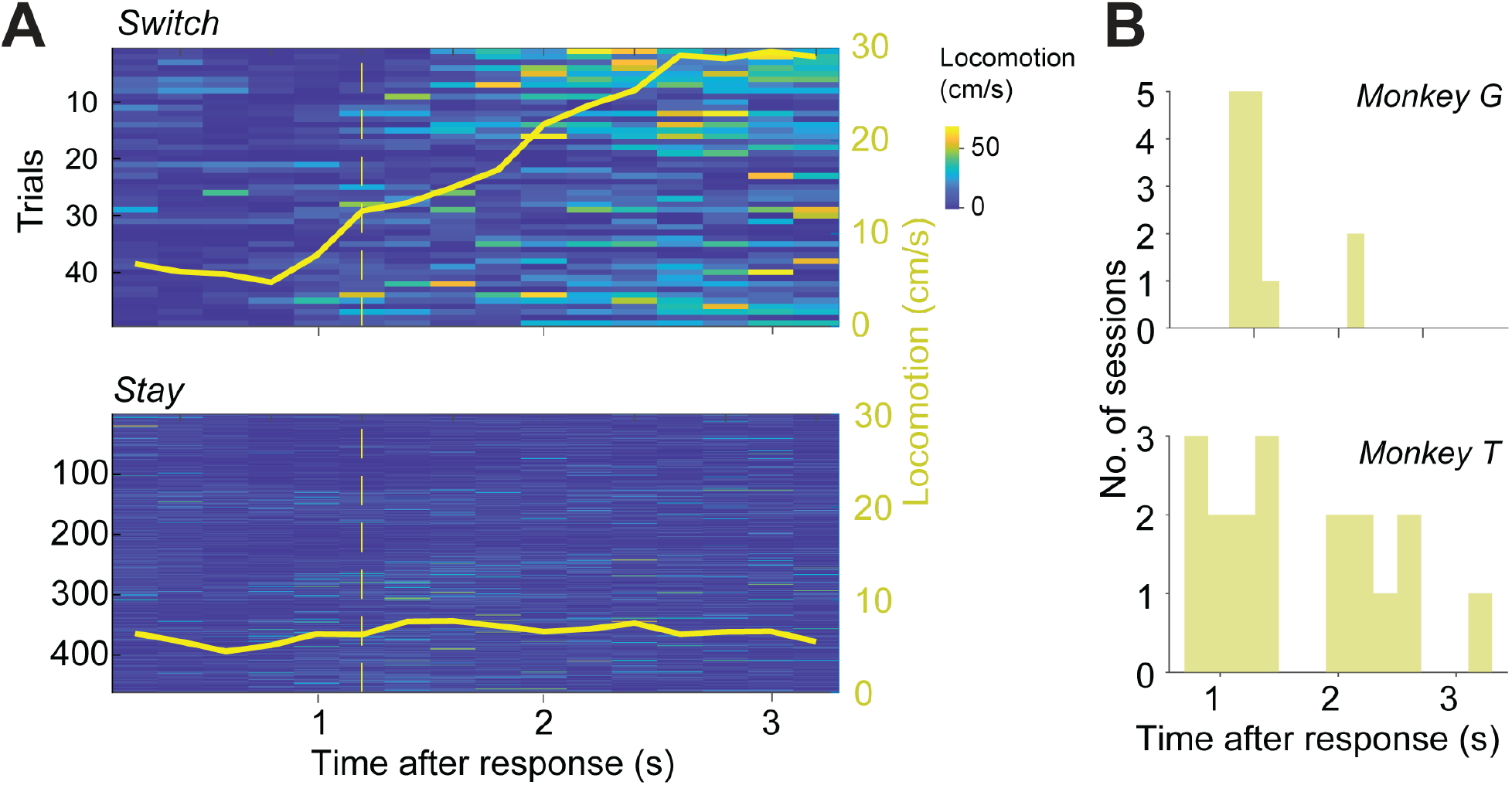
Detecting the time of the switches using the locomotion data. **(a)** the magnitude of locomotion in each 200 ms time bin, separated for switch and stay trials. The trial averages are shown as solid yellow lines. The yellow dashed line shows the time in which the average magnitude of locomotion in switch trials passes the 3 SD of the magnitude of locomotion in stay trials. **(b)** Average switch time in each session.

**Figure S9.**
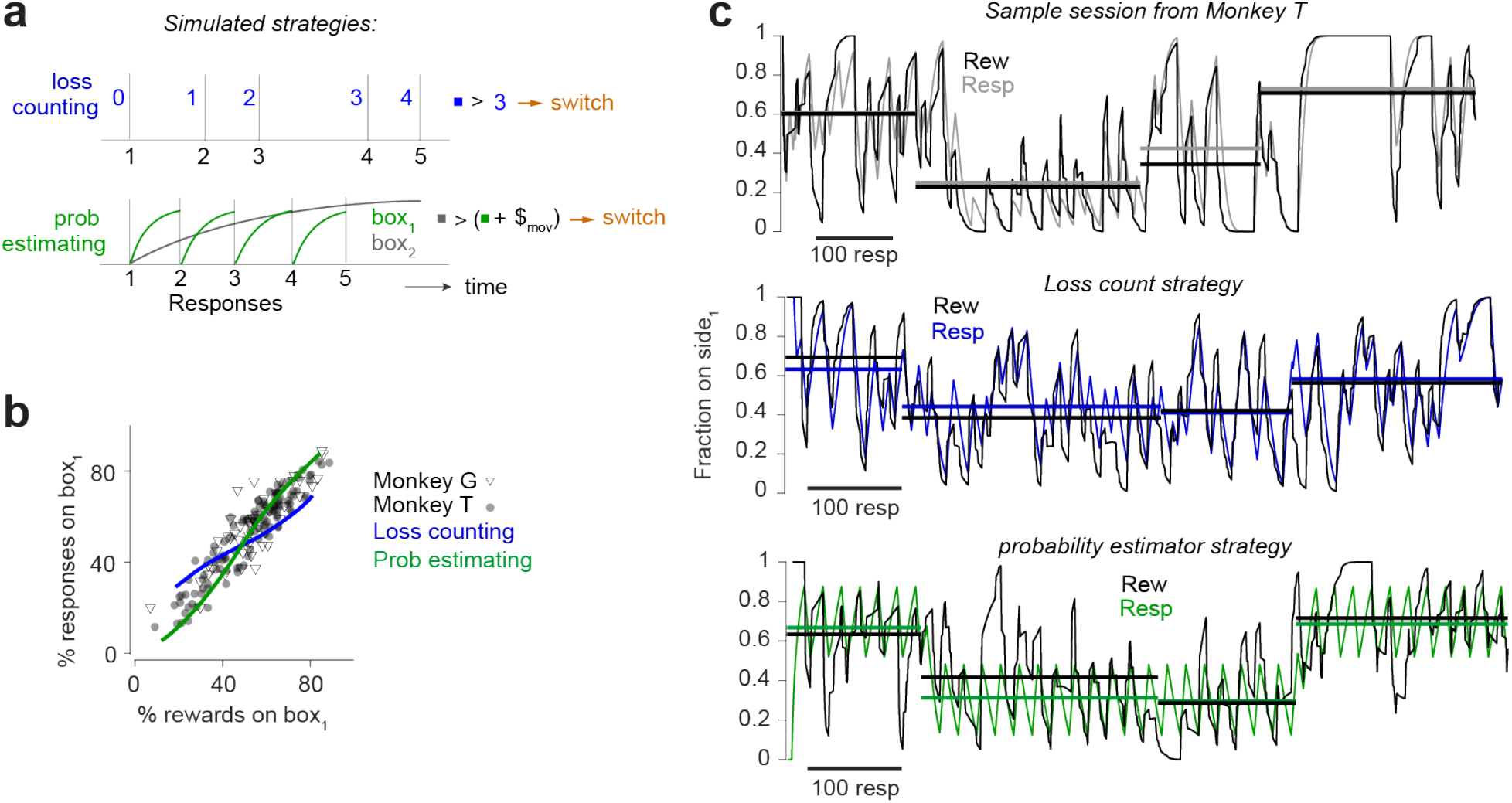
**(a)** Illustration of foraging strategies for two simulated agents. The ‘*loss counting*’ agent switches to the other box when *loss count* exceeds a threshold drawn from a Gaussian distribution (λ=2.66, σ=1.9). The ‘probability estimator’ agent switches to the other box when the probability of reward availability on the other box exceeds the probability of reward availability in the current box by a fixed switching cost ^1^. The inter-press times were drawn from a random geometric distribution for both agents. The parameters of these strategies, namely the *loss count* distribution, the inter-press time distribution, and the switching time, were estimated from the behavior of the monkeys. Each agent was simulated for 100 rewards for each set of reward schedules for box1 and box2. The variable interval (VI) reward schedules spanned the range between VI-5 and VI-50 in steps of 1 s and were drawn independently for each box. **(b)** Matching behavior of two monkeys and two simulated agents: the fraction of behavioral presses at box 1 is approximately proportional to the fraction of rewards at box 1 (26 sets of schedules for monkey G and 59 sets of schedules for monkey T are shown). Curves show spline fits for the matching behavior for the simulated agents. **c)** Dynamic matching for a sample session of Monkey T with 3 sets of reward schedules: VI15-VI25, VI25-VI15, and VI15-VI25 again. We compared two simulated agents of panel a. The reward and press rates were calculated locally using a causal Gaussian filter^6^.

